# Ketone supplementation dampens subjective and objective responses to alcohol in rats and humans

**DOI:** 10.1101/2023.09.23.558269

**Authors:** Xinyi Li, Zhenhao Shi, Dustin Todaro, Timothy Pond, Juliana Byanyima, Sianneh Vesslee, Rishika Reddy, Ravi Prakash Reddy Nanga, Gabriel Kass, Vijay Ramchandani, Henry R. Kranzler, Janaina C.M. Vendruscolo, Leandro F. Vendruscolo, Corinde E. Wiers

**Author notes:** **Correspondence:** Corinde E. Wiers, PhD, 3535 Market St., Suite 500, Philadelphia, PA 19104.

## Abstract

Previous preclinical and human studies have shown that high-fat ketogenic diet and ketone supplements (KS) are efficacious in reducing alcohol craving, alcohol consumption, and signs of alcohol withdrawal. However, the effects of KS on alcohol sensitivity are unknown. In this single-blind, cross-over study, 10 healthy participants (3 females) were administered a single, oral dose of a KS (25 g of ketones from D-β-hydroxybutyric acid and R-1,3-butanediol) or placebo 30 min prior to an oral alcohol dose (0.25 g/kg for women; 0.31 g/kg for men). Assessments of breath alcohol concentration (BrAC) and blood alcohol levels (BAL) and responses on the Drug Effect Questionnaire were repeatedly obtained over 180 min after alcohol consumption. In a parallel preclinical study, 8 Wistar rats (4 females) received an oral gavage of KS (0.42 g ketones/kg), water, or the sweetener allulose (0.58 g/kg) followed 15 min later by an oral alcohol dose (0.8 g/kg). BAL were monitored for 240 min after alcohol exposure. In humans, the intake of KS prior to alcohol significantly blunted BrAC and BAL, reduced ratings of alcohol liking and wanting, and increased disliking for alcohol. In rats, KS reduced BAL more than either allulose or water. In conclusion, KS altered physiological and subjective responses to alcohol in both humans and rats and the effects were likely not mediated by the sweetener allulose present in the KS drink. Therefore, KS could potentially reduce the intoxicating and rewarding effects of alcohol and thus be a novel intervention for treating alcohol use disorder.

## INTRODUCTION

The 2021 National Survey on Drug Use and Health estimated that approximately 21.5% of U.S. individuals aged 12 years or older binge drank alcohol in the past month, and that 10.2% of individuals met DSM-5 diagnostic criteria for alcohol use disorder (AUD) [1]. Excessive alcohol consumption increases the risk for reckless behaviors (e.g., violence and driving under the influence) and pathological conditions, including cognitive decline, liver disease, and certain types of cancer, and is one of the leading causes of preventable deaths in the United States [2].

Nutritional ketosis has emerged as a potential therapy for the treatment of alcohol withdrawal and AUD [3,4]. Nutritional ketosis is a metabolic state characterized by elevated levels of ketone bodies, which collectively refer to acetone, acetoacetate, and β-hydroxybutyrate. Ketosis can be achieved through fasting or adherence to a high-fat, low-carbohydrate ketogenic diet. Preclinical studies have shown the efficacy of a ketogenic diet in reducing alcohol consumption [4,5] and alcohol withdrawal [6,7]. These findings were corroborated by human studies demonstrating that a ketogenic dietary intervention attenuated alcohol craving and wanting [4,8] and the need for benzodiazepine medication for alcohol withdrawal management in AUD inpatients [4]. However, given the restrictive nature of the ketogenic diet, adherence has proven difficult. Exogenous ketone supplements (KS) offer an alternative and has been shown to elevate serum levels of ketone bodies without dietary deprivation of carbohydrates [9]. Preclinical studies have shown KS to reduce alcohol withdrawal symptoms and reinforcement similarly to a ketogenic diet [7]. However, the mechanisms underlying the efficacy of ketosis in reducing alcohol withdrawal, consumption, and craving/wanting are poorly understood and may involve physiological interactions with alcohol.

Alcohol is absorbed in the gastrointestinal tract, from which it enters the circulation for distribution throughout the body [10,11]. Alcohol is metabolized in the liver through sequential oxidation to acetaldehyde and acetate by alcohol and aldehyde dehydrogenases (ADH and ALDH), respectively. It is also metabolized to a lesser extent in non-hepatic organs, such as the stomach and the brain [10]. A preclinical study demonstrated five-fold blood alcohol levels following alcohol vapor exposure in rats that were maintained on a 6-week ketogenic diet compared to a standard chow diet [4]. These findings may be explained in part by the presence of ketones and/or the restriction of carbohydrates with a ketogenic diet, as both carbohydrate-rich diets [12] and sugar consumption [13,14] reduce breath alcohol concentration (BrAC) and blood alcohol levels (BAL) in responses to oral alcohol challenges. In contrast, oral exogenous KS in the presence of a regular, carbohydrate-rich diet significantly reduced BAL in mice compared to those maintained on a regular diet without KS or a ketogenic diet [7].

Here, we compared the acute effects of a KS with placebo on BrAC and BAL responses to an alcohol challenge in healthy human volunteers. We also assessed subjective responses to alcohol after KS and placebo on the Drug Effect Questionnaire, which includes questions regarding feelings of intoxication, alcohol liking, and wanting. Because the low-caloric sweetener allulose was present in our KS drink, we also compared the effects of KS with those of allulose on BAL responses to an oral alcohol challenge in rats. We hypothesized that, in contrast to a ketogenic diet [4], but consistent with previous KS findings [7], KS would reduce both objective (BrAC and BAL in both humans and rats) and subjective responses (in humans) to a moderate alcohol challenge.

## MATERIALS AND METHODS

### Human study

#### Participants

Twelve heathy individuals between the ages of 21 and 50 years were recruited for participation in the study. Participants reported to have consumed at least two standard alcohol drinks on a single day at least once in the prior month. Two participants withdrew from the study and n=10 participants (3 females and 7 males) completed both study arms. Participants were 29.7 ± 6.8 years of age, had a mean body mass index of 25.3 ± 4.6 kg/m^2^, and included one African American, one Asian, and eight Caucasians. All participants provided written informed consent and were paid for their participation. The study was approved by the Institutional Review Board at the University of Pennsylvania and registered at ClinicalTrials.gov (NCT05551754).

Participants were screened and excluded if they had any major medical condition (e.g., gastrointestinal disorders, including irritable bowel syndrome, liver disease, kidney disease, and metabolic disorders, including diabetes) or were taking medication that could interfere with study procedures (e.g., the use of ketone supplements or alcohol). Individuals with a major psychiatric or a substance use disorder (other than nicotine or cannabis use disorder) based on medical history and the structured Mini-International Neuropsychiatric interview for the Diagnostic and Statistical Manual of Mental Disorders, Fifth Edition [15] were excluded from participation. Also excluded from participation were females who were pregnant or lactating, individuals weighing greater than 225 lbs. (to limit the volume of alcohol provided to individuals), individuals positive for any substance other than cannabis on a urine drug test or who had a baseline breath alcohol level above 0.00% on the day of the study procedures.

#### Study design

In this single-blinded, cross-over study, participants received KS and placebo on two separate study visits with a minimum 1-day washout period in between. Participants were asked to arrive following an overnight fast and were provided a standardized meal (of approximately 550 kcal). Approximately 1 h later, participants drank the KS or placebo, followed by an oral dose of alcohol 30 min later (**Figure 1A**). All adverse events were mild and transient, and participants reported resolution of the side effects during follow-up phone calls the day after the study visit. A list of adverse events is provided in **Supplemental Adverse Events** and **Table S1**.

**Figure 1:**
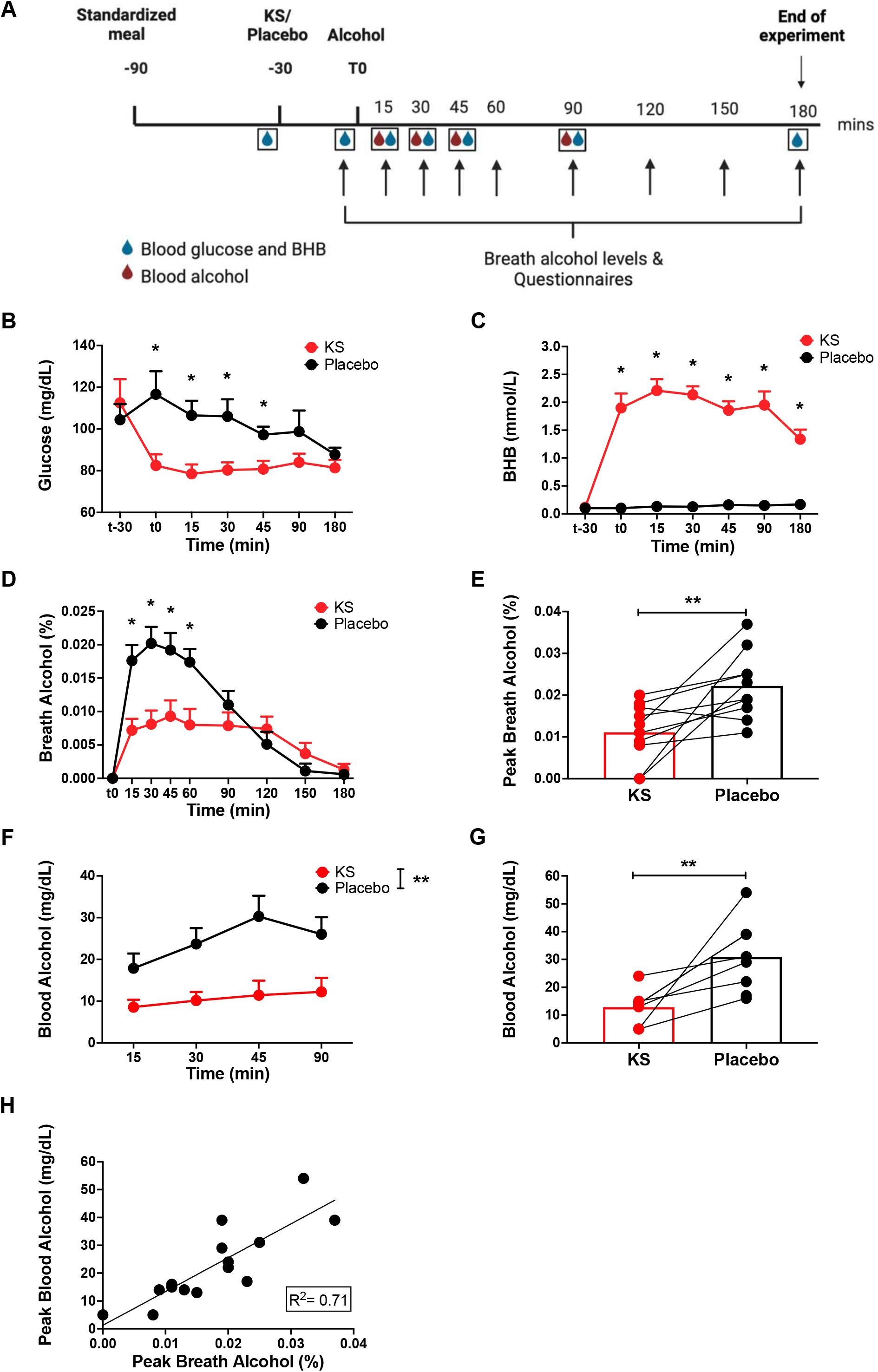
Responses following an alcohol challenge paradigm in individuals administered KS and placebo. **(A)**Schematic of the experimental timeline (human). Participants arrived at the study center following an overnight fast. They received a standardized meal, the KS or placebo approximately 1 h later and then an oral dose of alcohol 30 min later. BrAC (9 times, arrows), BAL (4 times, red drops), blood ketone and glucose levels (7 times, blue drops), and subjective ratings (5 times) were assessed. **(B)** Measures of blood glucose at t-30 (pre-intervention) (*n* =5/5; KS/placebo), t0 (pre-alcohol) (*n* =6/5), and at 15 (*n* =10/10), 30 (*n* =6/4), 45 (*n* =10/10), 90 (*n* =6/4), and 180 (*n* =10/9) min following alcohol administration. **(C)** Measures of blood BHB at t-30 (pre-intervention) (*n* =6/5; KS/placebo), t0 (pre-alcohol) (*n* =6/6), and at 15 (*n* =10/10), 30 (*n* =6/4), 45 (*n* =10/10), 90 (*n* =6/4), and 180 (*n* =9/8) min following alcohol administration. **(D** Measures of BrAC prior to (t0) (n=10/10; KS/placebo) and at 15, 30, 45, 60, 90, 120, 150, and 180 min (n=10/10) following alcohol administration**) (E)** Comparison of peak BrAC between placebo and KS. **(C)** Measures of BAL at 15 (*n* =8/7; placebo/KS), 30 (*n*=6/7), 45 (*n* =7/5), and 90 (*n* =6/5) min following alcohol administration. **(D)** Comparison of peak BAL between KS and placebo. **(E)** Correlation between peak BAL and BrAC. Mean ± SEM. **Indicates significant main Intervention effect. *Indicates significant differences between placebo and KS at a particular time point, *p* <0.05. Abbreviations: BHB, β-hydroxybutyrate; KS, ketone supplement.

#### Ketone supplement and placebo

The KS and placebo were supplied by Vitanav Inc, Washington DC. The KS (147.5 ml, Kenetik, Ketone Concentrate) contained 25 g of ketones from a mixture of D-β-hydroxybutyric acid and R-1,3-butanediol and 35 g of the low-calorie sweetener allulose (total 150 kcal). The placebo contained 74 ml of Safeway Sweet and Sour mix, 18.5 ml lemon juice, 18.5 ml lime juice, and 37 ml water (total 57.2 kcal from 14.3 g of sugar) (Vitanav Inc, Washington DC). The study drinks were diluted with water on the day of the study visit. The order in which the participants received the KS and placebo drinks was randomized and participants were blinded to the study intervention. After study participation, participants were asked which intervention they thought they received and rated their confidence in their guesses (0-10 scale).

#### Alcohol challenge

The dose of alcohol (100 proof Smirnoff vodka, Diageo plc, London, England) was adjusted for body weight and sex to achieve a target breath alcohol level of 0.05% (0.25 g/kg for women and 0.31 g/kg for men). The alcohol was diluted in sugar-free soda according to the participants’ preference and the participants were asked to finish the alcoholic drink within 5 min. Objective and subjective measures of alcohol intoxication were monitored prior to and for 180 min following alcohol intake. BrAC were assessed by handheld breath alcohol monitors (Intoximeters Inc., St. Louis, MO). Blood was collected by trained study staff and sent to the laboratory at the Hospital of the University of Pennsylvania for blood alcohol level (BAL) analyses. Due to difficulties in blood draws, several BALs were missed (Ns for each time points are presented in the Figure legends).

Self-report assessments included the Drug Effect Questionnaire [16] consisting of 5 items: (1) FEEL the effects of alcohol, (2) feel HIGH, (3) DISLIKE any of the effect you are feeling, (4) LIKE the effect you are feeling, (5) want MORE of what you consumed, which are rated on a 100-mm visual analog scale that ranges from Not at all (0) to A lot (100). The Brief Biphasic Alcohol Effect Scale (BAES) [17] assessed whether participants were Energized, Excited, Sedated, Sluggish, Up, or had Slow thoughts, on a 10-point scale from Not at all (0) to Extremely (10) . The responses were combined in the two sum scores: Stimulation (energized, excited, and up) and Sedation (sedated, slow thoughts, and sluggish). The 8-item Alcohol Urge Questionnaire (AUQ) [18] assessed alcohol craving. Responses on the DEQ and BAES were missing in one placebo participant at one and two time points, respectively.

Blood levels of ketone and glucose were collected before the KS/placebo intervention and monitored throughout the study using finger pricks and commercially available monitors (Precision Xtra, Abbott Laboratories, Chicago, IL) (Ns for each time points are presented in the Figure legends). Participants were provided a snack at the end of the study. In cases during which participants had blood glucose levels below 70 mg/dL (as was the case with two participants following the KS intervention trial), snacks were provided upon measurement of the blood sugar, which were monitored until they reached above 70 mg/ dL. Participants were called the next business day to inquire of any possible adverse events they may have experienced.

#### Data analyses

All analyses were conducted with SPSS (IBM Corp., Armonk, NY) using linear mixed-effects analyses to examine the main effect of Intervention (KS *vs*. placebo), main effect of Time (repeated assessments over time), and the Time x Intervention interaction effect. We included visit order (KS/Placebo or Placebo/KS) as a covariate and individual-specific random intercepts in the model. The study’s primary outcome measures were BrAC and BAL responses to KS *vs*. Placebo. For BAL, measurements below the laboratory detection cutoff of 10 mg/dL were analyzed as 5 mg/dL, as per Farokhnia et al. [19]. Secondary outcome measures were the effects of KS *vs*. placebo on the DEQ. Subjective responses to KS/placebo prior to alcohol intake were measured using the DEQ and compared using paired Student’s *t*-tests. To correct for potentially confounding effects of these pre-alcohol DEQ responses, we reported the linear mixed-effects analyses on the DEQ as both uncorrected and corrected for pre-alcohol ratings. When applicable, *post hoc* pairwise comparisons were analyzed with Bonferroni correction.

In exploratory analyses, we used linear regression to examine congruency between peak BAL and BrAC during the KS and placebo intervention arm, and linear mixed-effects analyses to examine the effects of the KS/placebo intervention on blood glucose and ketone levels, BAES and AUQ. Moreover, paired Student’s *t*-tests were utilized to compare peak BrAC and BAL following KS and placebo.

### Animal study

A total of 8 Wistar rats (4 males and 4 females) with ad libitum access to water and food were group-housed in a temperature- and humidity-controlled facility and maintained on a reverse 12-hour light/dark cycle (lights on at 6:00AM). All procedures were conducted according to the National Institutes of Health Guide for the Care and Use of Laboratory Animals and were approved by the National Institute on Drug Abuse, Intramural Research Program, Animal Care and Use Committee.

The rats received a dose of Kenetik (0.42 g ketones/kg), allulose (0.58 g/kg; Wholesome Sweeteners Inc., Sugar Land, TX), or water at a volume of 2.45 mL/kg of body weight through oral gavage. After 30 min, 0.8 g/kg of alcohol was administered orally. Blood was collected from the tip of the tail prior to and at pre-specified time intervals following alcohol administration (**Figure 3A**). Blood glucose (glucometer, Tyson Bioresearch Inc., Zhunan, Taiwan) and β-hydroxybutyrate (BHB; Precision Xtra Blood Ketone Monitoring System, Abbott, Alameda, CA), and alcohol levels were measured. Blood alcohol levels were assessed using gas chromatography-mass spectrometry, in which an ethanol calibration curve was prepared from 12.5 mg/dL to 300 mg/dL using ethanol standards in water (Cerilliant, Round Rock, TX). The 50 mg/dL calibrator was diluted with water to achieve calibrators at the 12.5 and 25 mg/dL levels. Briefly, 10 μL of ethanol standard or unknown whole blood was added to 10-mL glass headspace vials (Agilent, Santa Clara, CA) and sealed with a crimp cap. The vials were heated in a 70°C 7697A Headspace Sampler (Agilent) prior to headspace injection onto an MXT®-Volatiles 30 meter, 0.28 mm ID, 1.25 μm df column (Restek, Centre County, PA) using helium as the carrier gas. The 8890 GC System column oven (Agilent) was heated to 40°C for an isocratic 6-minute run paired with a 5977B GC/MSD (Agilent).

Blood glucose, BHB, and alcohol levels were analyzed using linear mixed-effects models with individual-specific random intercepts to test for the main effects of Intervention, Sex, and Time, as well as their interactions. If applicable, *post hoc* pairwise comparisons were analyzed with Bonferroni correction.

## RESULTS

### Human study

#### Blood glucose and BHB

For blood glucose levels, there were significant effects of Intervention (F_1,78.0_=27.6, *p* <0.001), Time (F_6,77.9_=5.7, *p* <0.001) and the Intervention x Time interaction (F_6,77.9_=5.7, *p* =0.008). *Post hoc* analyses showed significantly lower blood glucose levels for up to 75 min following KS *vs*. placebo administration (Bonferroni corrected, *p*<0.05) (**Figure 1B**).

For blood BHB, there were significant effects of Intervention (F_1, 76.0_=418.2, *p* <0.001), Time (F_6,75.4_=11.3, *p*<0.001), and the Intervention x Time interaction (F_6,75.4_=11.3, p<0.001). *Post hoc* analyses demonstrated significantly higher blood BHB levels following KS, which peaked at 45 min after KS intake and remained elevated throughout the 180-min alcohol challenge study, compared to placebo (Bonferroni corrected, *p* <0.05) (**Figure 1C**).

#### Breath and blood alcohol concentrations

For BrAC, there were significant effects of Intervention (F_1,152.0_=39.0, *p* <0.001), Time (F_8,152.0_=28.5, *p* <0.001), and the Intervention x Time interaction (F_8,152_=7.7, *p* <0.001) (**Figure 1D**). *Post hoc* analyses indicated significantly lower BrAC following KS than placebo for up to 60 min post-alcohol consumption (Bonferroni corrected, *p* <0.05). Furthermore, paired Student’s *t*-test demonstrated significantly lower peak BrAC with KS than placebo (t_9_=3.2, *p* =0.010) (**Figure 1E**). BrAC data for individual participants following KS and placebo are presented in **Supplemental Figure S1**.

For BAL, there was a significant effect of Intervention (F_1,37.3_=27.9, *p* <0.001), with lower BAL with KS. However, there were no effects of Time (F_3,37.0_=1.9, *p* =0.2) or the Intervention x Time interaction (F_3,37.3_=0.8, *p* =0.5) (**Figure 1F**). Lower peak BAL was observed following the KS intervention than placebo (t_6_=3.6, *p* =0.012) (**Figure 1G**). Exploratory linear regression with both KS and placebo measures pooled together demonstrated a significant association between peak BrAC and BAL measures (F_1,14_= 232.5, *p* <0.001, R^2^=0.71) (**Figure 1H**).

#### Subjective ratings – DEQ

At t0 (i.e., 15 min after KS/Placebo but prior to alcohol administration), participants reported lower wanting MORE (t_9_=3.0, *p* =0.015) and higher DISLIKE at a trend level (t_8_=-2.17, *p* =0.06) for KS *vs*. placebo. There were no effects of Intervention on HIGH, LIKE, or FEEL the effects of alcohol at t0 (**Figure 2**).

**Figure 2:**
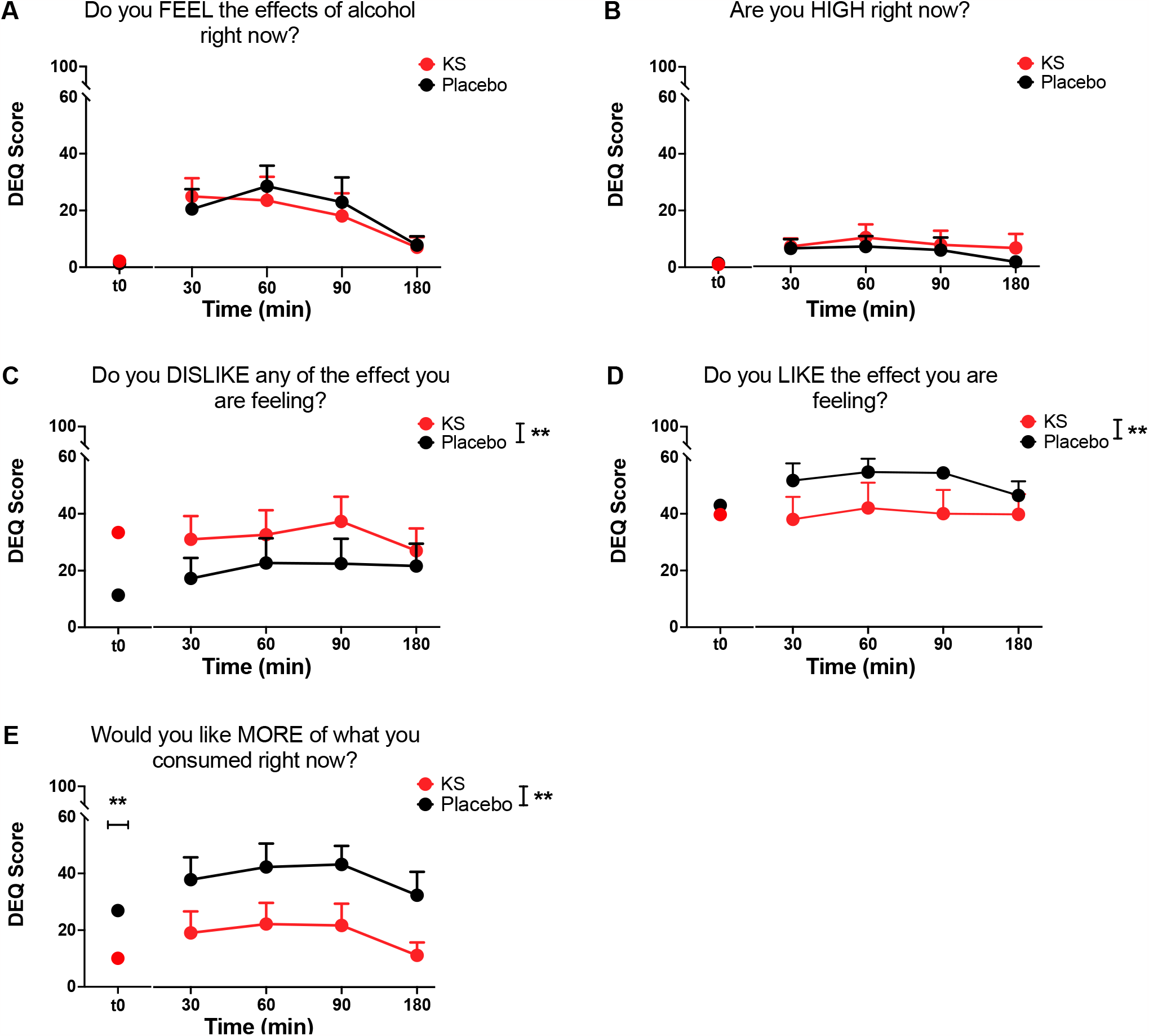
Drug Effect Questionnaire responses before and following an alcohol challenge paradigm in individuals administered KS or placebo. **(A)** Subjective responses regarding **(A)** FEEL the effects of alcohol (**B**) reports of HIGH, **(C)** DISLIKE the effects, **(D)** LIKE the effects, **(E)** like MORE of what you consumed assessed prior to and at 30, 60, 90, and 180 min following the alcohol challenge. Mean ± SEM. **Indicates significant main effect of Intervention, unadjusted, *p* <0.05. Abbreviations: DEQ, Drug Effect Questionnaire; KS, ketone supplement.

After alcohol intake, there were significant main effects of the KS Intervention on DEQ ratings of DISLIKE (F_1,57.1_=9.4, *p* =0.003), LIKE (F_1,56.6_=6.8, *p* =0.012) and liking MORE (F_1,60.3_=29.8, *p* <0.001), with KS resulting in higher DISLIKE, and lower LIKE and wanting MORE ratings of alcohol. There were no effects of the Intervention on FEEL the effects of alcohol (F_1,60.3_= 0.7, *p* =0.4;) or HIGH (F_1,60.3_= 0.8, *p* =0.4) **(Figure 2**). There was a significant main effect of Time on FEEL the effects of alcohol (F_3,60.2_= 5.8, *p* =0.002) but not on the other DEQ items (*p*s >0.05). There was no significant Intervention x Time interaction (*p*s >0.05). The significant main effects of KS Intervention remained after correction for pre-alcohol ratings for DISLIKE (F_1,58.8_=8.8, *p* =0.004) and LIKE (F_1,53.4_=5.7, p=0.02), but not for wanting MORE alcohol (F_1,65.6_=0.0, p=0.9) (See **Supplemental Analyses of Baseline-Adjusted DEQ scores** for analyses that correct for pre-alcohol ratings).

#### Subjective ratings – BAES and AUQ

There were no significant effects of Intervention, Time, or the Intervention x Time interaction on BAES or AUQ ratings (all *p*s >0.05, see **Supplementary Analyses of BAES and AUQ** and **Supplemental Figure S2**).

#### Study blinding

Eight of 10 participants during the KS and nine of 10 participants during the placebo visit correctly guessed the study intervention (χ_1_=9.9, *p* =0.002). Confidence ratings did not differ between the KS (mean= 5.6 ± 3.1) and Placebo (5.8 ± 3.0) visits (t_9_=0.3, *p* =0.8).

### Animal study

For blood glucose levels, there were significant effects of Time (F_5, 90_=8.4, *p* <0.001) and Sex (F_1, 18_=45.2, *p* <0.001), but no main effect of Intervention (F_2, 18_=0.0, *p* =1.0) or any interaction effects (*p*s >0.05) (**Figure 3B**). For blood BHB levels, there were significant effects of Intervention (F_2, 18_=17.7, *p* <0.001), Time (F_5, 90_=4.0, *p* =0.002), and the Intervention x Time interaction (F_10, 90_=2.4, *p* =0.014). *Post hoc* comparisons indicated that blood BHB was significantly higher up to 150 min after KS intake than water and allulose (Bonferroni corrected, *p* < 0.05) (**Figure 3C**). There were also significant effects of Sex (F_1, 18_=7.5, *p* =0.014) and the Time x Sex interaction (F_5, 90_=2.4, *p* =0.046), such that males had higher BHB levels than females at 30, 60, and 240 min following alcohol consumption (Bonferroni corrected, *p* <0.05).

**Figure 3:**
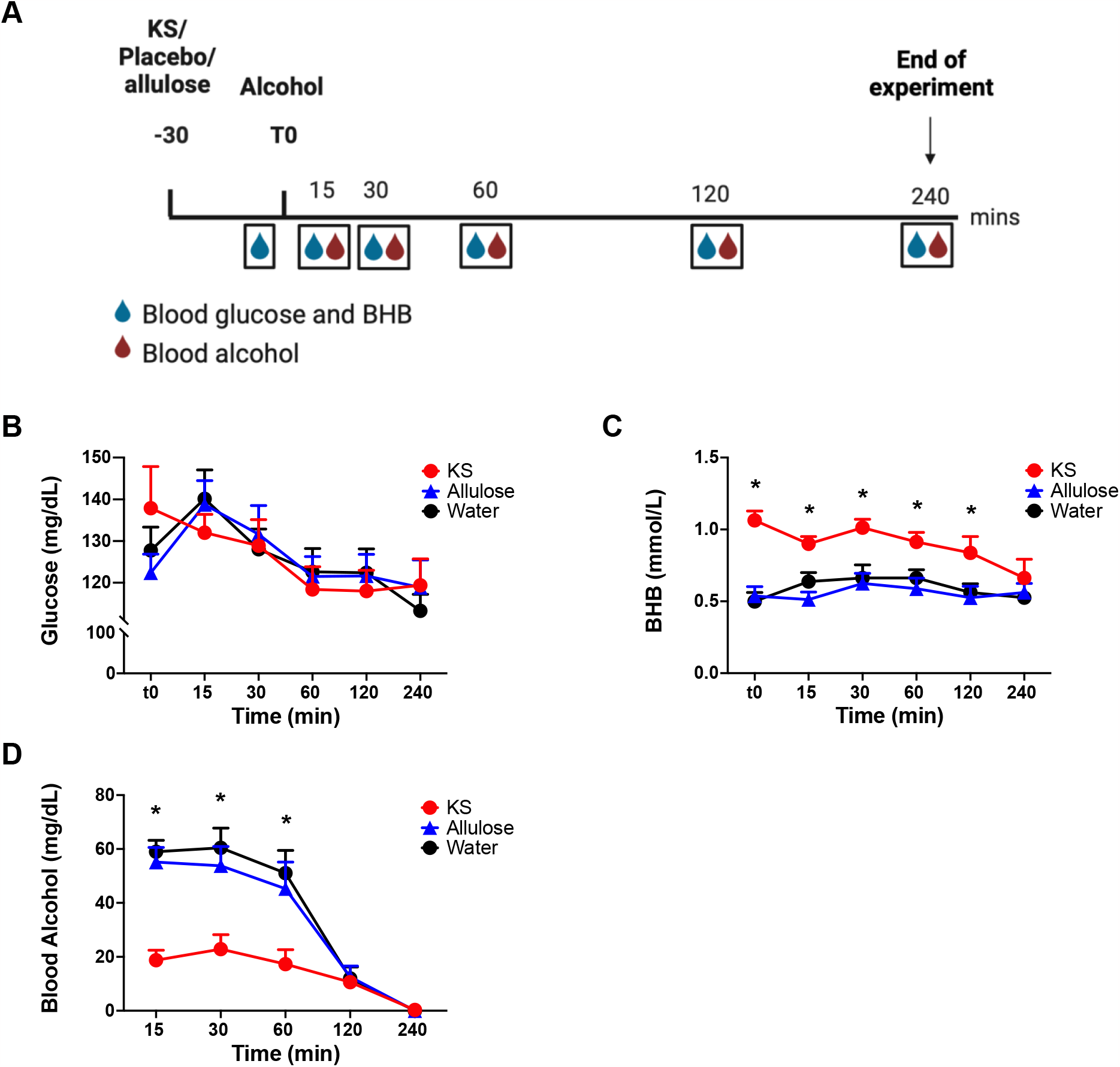
Responses levels following an oral alcohol challenge in rats administered KS, water, or allulose. **(A)** Schematic of experimental timeline (rats). Rats were administered KS, allulose, or water orally 30 min prior to an oral dose of alcohol. Blood was collected for assessments of blood alcohol levels (5 times, red drops) and blood ketone and glucose levels (6 times, also collected at t0; blue drops). **(B)** Blood glucose and **(C)** BHB levels were assessed immediately prior and at 15, 30, 60, 120, and 240 min post-alcohol challenge. **(D)** Assessments for BAL were made at 15, 30, 60, 120, and 240 min after oral administration of alcohol. Mean ± SEM. *Indicates significant KS differences from water and allulose at a particular time point, *p* <0.05. Abbreviation: BHB, β-hydroxybutyrate; KS, ketone supplement.

Analyses of BAL in rats indicated a significant effect of Intervention (F_2,20.4_=21.9, p<0.001), Time (F_4,70.9_=119.2, p<0.001), Sex (F_1,16.2_=14.1, *p* =0.002), the Intervention x Time interaction (F_8, 70.0_=11.0, p<0.001), and the Sex x Time intervention (F_4,70.9_=8.6, *p* <0.001), but not the Intervention x Time x Sex interaction (F_10,46.5_=0.8, *p* =0.6) (**Figure 3D**). *Post hoc* pairwise comparisons did not show significant differences in BAL between water and allulose. KS significantly reduced BALs compared to water and allulose up to 60 min following alcohol consumption (Bonferroni corrected, *p* <0.05). Male mice demonstrated higher BAL than females at 30, 60, and 120 min following alcohol administration (Bonferroni corrected, *p* <0.05).

## DISCUSSION

Emerging studies have shown a role for nutritional ketosis, induced through a ketogenic diet or exogenous KS, in alleviating symptoms of alcohol withdrawal and alcohol craving [3,4,7]. Here, we found evidence in both humans and rats that the administration of a KS prior to alcohol significantly attenuated breath alcohol concentration and blood alcohol levels. In humans, KS additionally reduced subjective ratings of alcohol liking and wanting, and increased alcohol disliking. However, no differences in reports of feeling high, feeling the effects of alcohol, or feeling stimulated or sedated by alcohol were observed in participants following KS *vs*. placebo. The lack of effects on subjective ratings of intoxication may be due to the low dose of alcohol that was administered (i.e., mean BrAC level of 0.02 in the placebo condition). Nevertheless, our findings suggest that KS modulates alcohol bioavailability and reduces the rewarding effects of alcohol (i.e., reduced alcohol liking and wanting). Because the KS used in the study contained high quantities of the sweetener allulose, which can be synthetically converted from allitol by alcohol dehydrogenase [20], a key enzyme in alcohol metabolism, we also evaluated the potential confound that allulose alone may have on blood alcohol levels in rats. We found that allulose had no effects on BAL in rats, whereas the KS decreased BAL, which further corroborated our findings that KS modulate alcohol bioavailability.

We postulate the following mechanisms by which KS reduces alcohol bioavailability. First, ketone D/L-3-Hydroxybutyrate has been shown to delay gastric emptying in healthy humans [21], and, delayed gastric emptying is associated with lower alcohol absorption [22]. Second, studies have shown that exogenous ketones increase levels of nicotinamide adenine dinucleotide (NAD+) [23,24], a cofactor necessary for alcohol metabolism [11], and increased NAD availability may accelerate alcohol metabolism in the liver [25]. Third, 1,3-butanediol is catabolized to BHB by ADH [26] and may prime the activity of ADH for the subsequent metabolism of alcohol. In one study, in a sub-population of individuals, prior exposure to alcohol increased the elimination rate of a second dose of alcohol by more than 40% above the basal rate [27]. Therefore, a potential prior activation of ADH, either by alcohol or 1,3 butanediol, could increase the metabolic rate of alcohol. Further research is needed to elucidate the mechanisms underlying the effects of KS and to differentiate effects on alcohol absorption from those on metabolism.

Our finding that KS *reduced* BrAC and BAL in humans and rats contrasts with our previous findings that endogenous production of ketones following a 6-week ketogenic diet substantially increased blood alcohol levels in rats exposed to alcohol vapor [4]. These results may suggest that the KS slows down alcohol absorption rather than speeding up alcohol metabolism because alcohol administered through inhalation is absorbed mainly through the lungs, bypassing gastrointestinal absorption and first pass metabolism in the stomach, and rapidly reaches the arterial circulation [28,29]. Consistent with the present study results, oral administration of KS and alcohol in mice also led to significantly dampened BAL [7]. However, Bornebusch et al. also indicated that a 3-week ketogenic *vs*. regular diet had no effect on BAL following oral alcohol administration [7]. The difference in findings with the ketogenic diet and KS may also be due to the carbohydrate restriction required by the diet. Thus, it is possible that the carbohydrate-restriction characteristic of the ketogenic diet, rather than the presence of ketone bodies *per se*, impaired alcohol metabolism in the rats. There is evidence that carbohydrates [12], including sucrose and fructose [13,14], reduce BrAC and BAL, due to both delayed absorption and increased availability of NAD+ for alcohol metabolism. Here, KS, without adherence to a carbohydrate-restrictive diet, elevated ketone levels. Moreover, studies have shown a greater ADH concentration in the liver of rats in the fed than the fasting state [30,31]. Administration of a 6-week ketogenic diet could induce a fasting state resulting in increased ADH activity and a decreased rate of alcohol oxidation.

Previous clinical and preclinical studies have shown that a ketogenic diet decreases alcohol craving and alcohol consumption [4,5]. Although, here, the KS reduced alcohol liking and wanting ratings, we did not see significant effects of KS on alcohol craving scores as measured with the AUQ. The null finding is likely attributable to the inclusion of healthy individuals with low alcohol craving scores. Future studies are needed to examine the effects of KS on alcohol craving and alcohol consumption in individuals with AUD and in preclinical models of alcohol dependence. Interestingly, we did not find significant effects of the KS on subjective feelings of intoxication. The low dose of alcohol used in this study may not have been sufficient to elicit feelings of intoxication, as the BAES responses during the alcohol challenge did not change over time. Furthermore, exploratory analyses did not show significant correlations between BrAC/BAL and subjective ratings of intoxication on the BAES or DEQ (**Supplemental Table S2**). The presence of sucrose in our placebo drink may also have dampened the intoxicating effects of alcohol [13,14] and contributed to the absence of differences between KS and placebo. Future studies should examine the effects of KS on both subjective and objective responses to a higher dose of alcohol.

Consistent with Stubbs, et al. ^9^, we demonstrated that a single dose of a KS was effective in elevating blood BHB levels and decreasing blood glucose within 30 min of consumption. Several incidences of hypoglycemia (glucose <70 mg/dL) were observed during the study that may be attributable to the KS alone or the interaction of KS with alcohol. Stubbs et al. (2017) found that KS increased insulin levels, which could account for the reduced glucose levels. In their seminal paper, Krebs et al. (1969) found that alcohol inhibits gluconeogenesis, which produces glucose from non-carbohydrate carbons (i.e., lipids and proteins), and these findings were later replicated in human studies that measured gluconeogenic flux following radiolabeled [2-^13^C_1_]glycerol infusion [32]. Although suppression of gluconeogenesis may not be concerning under normal physiological conditions when glucose is readily available, the effect may be compounded by alcohol consumption combined with KS, which together may give rise to hypoglycemia. None of our participants reported symptoms of hypoglycemia (e.g., dizziness and shakiness), likely because the high levels of ketones provided an alternative to glucose as a source of energy. Nonetheless, the effect of KS on glucose levels raises potential health concerns, for example in individuals with metabolic disorders such as diabetes. Furthermore, KS-associated gastrointestinal discomfort was reported in our study and others [33], but the effects were mild and transitory, and may be attributable to the ketones, allulose, or both.

Our study has several limitations and raises questions for future studies. First, our sample of healthy individuals was small (*n* =10) and future larger studies that include individuals with AUD are warranted to compare the effects of KS on alcohol-related measures in healthy and AUD individuals. Furthermore, low alcohol craving scores in our study population of healthy individuals likely resulted in a floor effect that obscured any effects of KS on alcohol craving. Second, we chose a low alcohol dose due to concerns about heightened intoxication in individuals following KS, as was observed in rats on a ketogenic diet [4]. The low dose of alcohol may have been insufficient to elicit intoxicating effects, as exploratory analyses yielded no correlation between BrAC/BAL and BAES and DEQ scores (**Supplemental Table S2**). This could account for the null effects of KS on subjective reports of intoxication. Third, our placebo and KS were not calorically matched, which could have contributed to the observed differences in alcohol sensitivity. However, our placebo contained sucrose, which has been shown to reduce alcohol sensitivity [13,14]. We found that KS dampened BrAC and BAL more than a sugar-containing placebo, which testifies to the robustness of the effects of KS. Fourth, the mechanism(s) underlying the effects of KS on alcohol absorption and metabolism remain to be elucidated. In our study, compared to placebo, KS reduced peak BrAC and BAL, but exploratory data analyses revealed no difference in elimination rates [t(7)=1.4, *p* =0.2] (**Supplemental Analyses of Alcohol Elimination Rate** and **Figure S4**). This may reflect an effect of KS in reducing alcohol absorption, rather than altering metabolism [34], but future studies are warranted to differentiate the two mechanisms. Fifth, the taste of KS was difficult to conceal, thus most participants correctly identified the KS and placebo interventions. Although we controlled for baseline measures (t0, post-KS/placebo intervention and pre-alcohol administration), it is possible that the unpleasant taste and nauseating effects of KS could have confounded subjective responses to alcohol.

In sum, we found that the administration of an exogenous KS attenuates breath alcohol concentration and blood alcohol levels, increases “disliking”, and decreases “liking” and “wanting” alcohol. Binge alcohol drinking and AUD are prevalent in the United States [1] and alcohol intoxication increases the risks for alcohol poisoning and reckless behaviors that are harmful to oneself or others [35]. Therefore, KS may have therapeutic potential to reduce alcohol intoxication and alcohol-related harm by reducing blood alcohol levels.

## AUTHOR CONTRIBUTION

XL, TP, RPRN, HRK JCMV, LFV, CEW contributed to study conceptualization and design. JB, SV, RR, GK were responsible for participant recruitment and data collection. ZS, GK, VR, HRK, JCMV, LFV, and CEW analyzed and interpreted the data. XL, ZS, DT, and CEW drafted the manuscript and all authors provided edits to the manuscript.

