## Supplemental Material for "Ketone supplementation dampens subjective and objective responses to alcohol in rats and humans"

### SUPPLEMENTARY MATERIALS

**Table S1**. Summary of adverse events.

| Number of participants | Event | Severity | Relatedness to KS |
|---|---|---|---|
| 3 (including 1 drop out) | Gastrointestinal discomfort: nausea, stomachache, diarrhea, and vomiting (drop-out) | Mild | Probable |
| 1 | Fainting during blood draw | Mild | Unlikely |
| 5 | Glucose < 70 mg/ dl during at least 1 time point during the study | Mild | Probable |

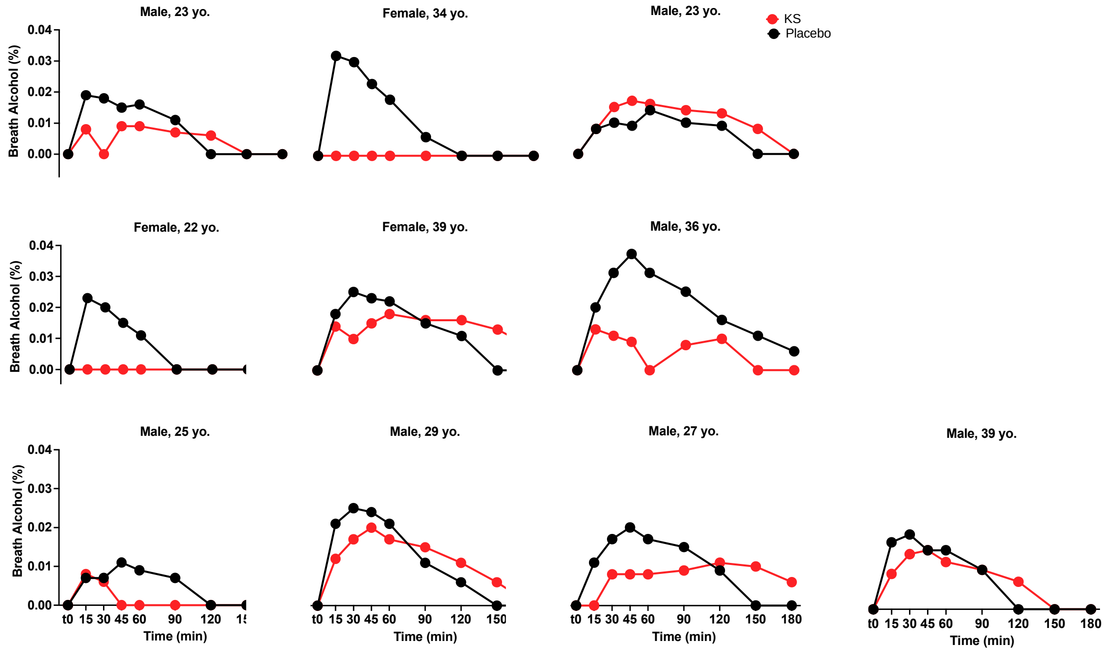


**Figure S1**. **Individual breath alcohol concentration following an alcohol challenge in individuals administered KS or placebo.** BrAC was measured prior to (t0) and 15, 30, 45, 60, 90, 120, 150, and 180 min following following alcohol administration. Abbreviations: KS, ketone supplement.

### Analyses of Baseline-Adjusted DEQ Scores

For "FEEL(ING) the effects of alcohol", there was significant effect of Time ($F_{3,59.3}$= 5.7, $p$ =0.002), but not Intervention ($F_{1,60.7}$= 0.6, $p$ =0.4) or Time x Intervention ($F_{3,59.3}$= 0.4, $p$ =0.7). Reports of "HIGH" throughout the alcohol challenge did not differ with effects of Intervention ($F_{1,63.4}$= 1.0, $p$ =0.3), Time ($F_{3,60.0}$= 0.9, $p$ =0.4), or Time x Intervention ($F_{3,60.1}$= 0.3, $p$ =0.8). For "DISLIK(ING) any of the effects you are feeling", there was a significant effect of Intervention ($F_{1,58.8}$=8.8, $p$ =0.004), but no effects of Time ($F_{3,52.8}$= 0.7, $p$ =0.5) or Intervention x Time interaction ($F_{3,52.7}$= 0.2, $p$ =0.9). Reports of "LIK(ING) any of the effects you are feeling" demonstrated an effect of Intervention ($F_{1,53.4}$=5.7, $p$ =0.02), but no significant effects of Time ($F_{3,53.2}$= 0.3, $p$ =0.8) or Intervention x Time interaction ($F_{3,53.2}$= 0.2, $p$ =0.9) were observed. For responses to "lik(ing) MORE of what you consumed", there was a significant effect of Time ($F_{3,57.9}$= 3.0, $p$ =0.039), but not Intervention ($F_{1,65.6}$=0.0, $p$ =0.9) or Intervention x Time interaction ($F_{3,57.9}$= 0.1, $p$ =1.0).

**<u>Analyses of BAES and AUQ</u>**

On the BAES, paired t-test did not reveal significant baseline (t0) differences on subjective reports of stimulation ["energized," "excited," and "up", t(9)=1.2, $p$ =0.2) or sedation ["sedated", "slow thoughts", and "sluggish", t(9)=1.6, $p$ =0.2). Reports of stimulation did not differ with effects of Intervention ($F_{1,61.2}$=3.4, $p$ =0.07), Time ($F_{3,61.1}$=0.2 $p$ =0.9) or Intervention x Time interaction ($F_{3,61.1}$=31.0 p=0.4), even after adjusting for baseline responses [Intervention: ($F_{1,66.4}$=0.001, $p$ =0.98); Time: ($F_{3,60.8}$=0.3, $p$ =0.8); Intervention x Time interaction ($F_{3,60.8}$=1.3 $p$ =0.3)]. Reports of sedation did not significantly differ with effects of Intervention ($F_{1,61.1}$= 1.6, p=0.2), time ($F_{3,61.0}$= 0.5, p=0.7), or Intervention x Time interaction ($F_{3,61.0}$= 0.4, p=0.8) and no significant effects were observed after adjusting for baseline responses [Intervention ($F_{1,64.3}$= 1.4, $p$ =0.2); Time ($F_{3,59.9}$= 0.7, $p$ =0.6); Intervention x Time interaction ($F_{3,59.9}$= 0.6, $p$ =0.6) (see **Figure S3A**).

Regarding participants' responses on the AUQ, no significant effects of Intervention ($F_{1,44.0}$= 0.2, $p$ =0.7), Time ($F_{2,44.0}$= 0.4, $p$ =0.7), and Intervention x Time interaction ($F_{2,44.0}$= 0.5, $p$ =0.6) were observed (see **Figure S3B**).

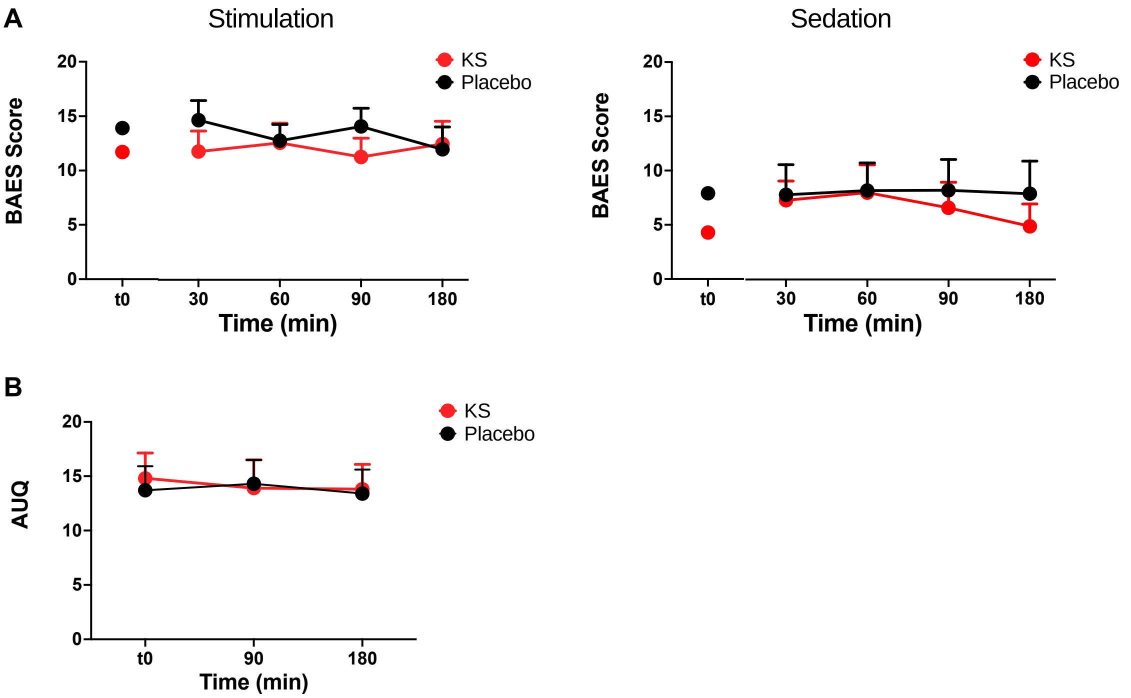


**Figure S2**. **Subjective responses following an alcohol challenge in individuals administered KS and placebo.** (A) BAES responses before and 30, 60, 90, and 180 min following an alcohol challenge. (B) AUQ responses before and 90 and 180 min after an alcohol challenge. Mean ± SEM. Abbreviations: AUQ, alcohol urge questionnaire; BAES, brief biphasic alcohol effect scale; KS, ketone supplement.

**Table S2**. Correlation between breath and blood alcohol levels with subjective report of intoxication.

| Questionnaire items | Breath alcohol (%) | Blood alcohol (mg/dL) |
|---|---|---|
| ***BAES*** | | |

| | | |
|---|---|---|
| Stimulation | $R^2$=0.1, $p$ =0.2 | $R^2$=0.04, $p$ =0.3 |
| Sedation | $R^2$=0.05, $p$ =0.8 | $R^2$=0.01, $p$ =0.7 |
| ***DEQ*** | | |
| Do you FEEL the effects of alcohol right now? | $R^2$=0.06, $p$ =0.3 | $R^2$=0.09, $p$ =0.3 |
| Are you High, right now? | $R^2$=0.00, $p$ =1.0 | $R^2$=0.01, $p$ =0.7 |
| Do you DISLIKE any of the effects you are feeling? | $R^2$=0.01, $p$ =0.6 | $R^2$=0.03, $p$ =0.6 |
| Do you LIKE the effects you are feeling? | $R^2$=0.01, $p$ =0.6 | $R^2$=0.06, $p$ =0.4 |
| Would you like MORE of what you consumed right now? | $R^2$=0.2, $p$ =0.08 | $R^2$=0.1, $p$ =0.2 |

### Analyses of Alcohol Elimination Rate

Alcohol elimination rates were calculated as per Jones, 2019. Individual Pearson correlation coefficients were extracted from linear regression between time and BrAC in the descending phase of the alcohol curve (i.e., following peak BrAC) and compared between KS and placebo intervention arms using paired t-test. We were unable to perform linear regression and calculate correlation coefficients for 2 participants during the KS intervention because they had BrAC of 0 throughout the alcohol challenge. We did not find a significant difference in elimination rate between the KS and placebo interventions ($t_7$=1.4, $p$ =0.2) (see **Figure S1**).

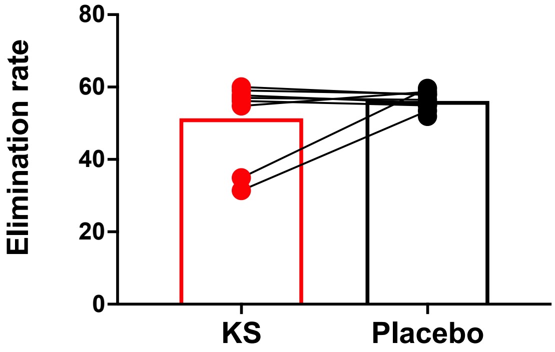


**Figure S3. Alcohol elimination rate following an alcohol challenge paradigm in individuals administered KS or placebo administration.** Abbreviations: KS, ketone supplement.
